# dBBQs : dataBase of Bacterial Quality scores

**DOI:** 10.1101/187641

**Authors:** Visanu Wanchai, Preecha Patumcharoenpol, Intawat Nookaew, David Ussery

## Abstract

**Background:** It is well-known that genome sequencing technologies are becoming significantly cheaper and faster. As a result of this, the exponential growth in sequencing data in public databases allows us to explore ever growing large collections of genome sequences. However, it is less known that the majority of available sequenced genome sequences in public databases are not complete, drafts of varying qualities. We have calculated quality scores for around 100,000 bacterial genomes from all major genome repositories and put them in a fast and easy-to-use database.

**Results:** Prokaryotic genomic data from all sources were collected and combined to make a non-redundant set of bacterial genomes. The genome quality score for each was calculated by four different measurements: assembly quality, number of rRNA and tRNA genes, and the occurrence of conserved functional domains. The dataBase of Bacterial Quality scores (dBBQs) was designed to store and retrieve quality scores. It offers fast searching and download features which the result can be used for further analysis. In addition, the search results are shown in interactive JavaScript chart framework using DC.js. The analysis of quality scores across major public genome databases find that around 68% of the genomes are of acceptable quality for many uses.

**Conclusions:** dBBQs (available at http://arc-gem.uams.edu/dbbqs) provides genome quality scores for all available prokaryotic genome sequences with a user-friendly Web-interface. These scores can be used as cut-offs to get a high-quality set of genomes for testing bioinformatics tools or improving the analysis. Moreover, all data of the four measurements that were combined to make the quality score for each genome, which can potentially be used for further analysis. dBBQs will be updated regularly and is freely use for non-commercial purpose.

## Background

It is well known that the current state-of-art of sequencing technologies makes genome sequencing significantly cheaper and quicker. Especially, the third generation sequencing which based on single-molecule sequencing technologies, have gained popularity because of ability of generating the long read [1]. Also, the exponential growth in sequencing data in public databases allow us to explore through large collections of genome sequences [2]. However, it is less known that many genomes in public databases are left as draft genome sequences. A huge number of draft genomes usually comes from difficulty of finishing process of genome sequences generated by second generation sequencing machine. Therefore, many genome projects on major genome repositories were left unfinished [3].

The estimation of errors in draft genome by Denton et al. [4] in 2014 indicated that, by comparing the same genomes with different level of completeness, nearly 40% of all gene families were inferred to have incorrect number of genes in draft genomes. Also, the possible reason of having over predicted genes in unfinished genomes is the fragmentation of genes in many contigs. Hence, these non-finished genome sequences may vary in qualities causing the inconsistent analysis.

Here, we collected both draft and complete genomes for around 100,000 bacterial genomes from major genome repositories: GenBank and GenBank Sequence Read Archive provided by the National Center for Biotechnology Information [5], the Broad Institute [6], the U.S. Department of Energy Systems Biology Knowledgebase, and the Pathosystems Resource Integration Center [7]. Then all genomes were annotated and assessed for the quality scores with the same method. We designed and implemented database to stores all genomes and their analysis. The website was constructed by the concept of interactive designed which allows users to interact directly with data and get feedback instantly.

## Construction and content

### Data sources

We retrieved bacterial genomic data from 4 different sources: GenBank, GenBank - SRA, Broad, Kbase, and PATRIC. These databases are major public genome repositories containing all types of genome completeness ranging from complete gnomes to contigs. The detail of retrieving genome sequences for each database can be described as follows. GenBank genomes were retrieved from the FTP site provided by NCBI [8]. Then each of whole genome sequence in Fasta nucleotide format was download and stored in a directory. The Fasta files of GenBank - SRA, which have already assembled as previously reported by the work of Larsen et al. [9], were downloaded from NCBI SRA FTP site [10] and stored the same way that previously described with GenBank genomes. The genome data from Broad Institute were retrieved from the Broad Olive website [6]. The bundle files of Broad project were extracted and kept only Fasta files. Kbase genomes were obtained through its API which allowed us to easily select any level of completeness and only Fasta format for genomes. Fasta files from PATRIC were searched and downloaded by using its FTP site [11].

### Genome quality scores

The genome quality score for each genome was calculated using the method proposed by Land et al. [12]. In order to standardize all genomes in the analysis, all genomes in Fasta format from different sources were predicted for the protein-coding genes using Prodigal [13]. Next, all 4 individual scores—Sequence Quality score, rRNA score, tRNA score, and Essential gene score—were calculated using RNAmmer [14], tRNAscan-SE [15], and HMMER3 [16] with Pfam-A [17] respectively (Figure 1). Basically, each of score is the measurement of the completeness of genome sequence: assembly quality, number of rRNA and tRNA genes, and the presence of conserved functional domains. Then all 4 scores were averaged to estimate the genome quality score.

**Figure 1.**
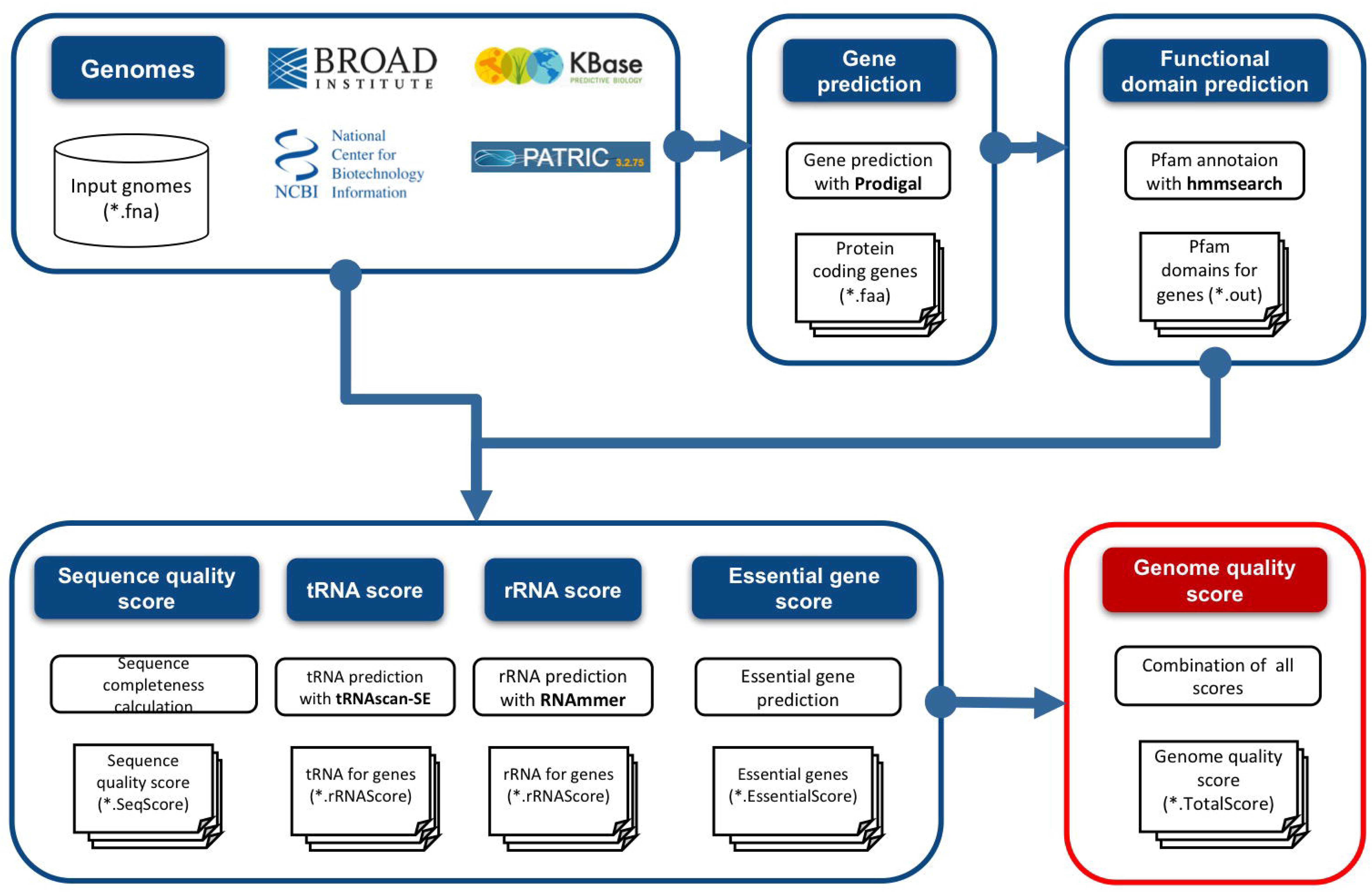
Analysis workflow of genome quality scores.

### Database schema and implementation

The database of dBBQs was developed as a relational database using SQlite3. The Apache HTTP 2.4.6 web server was then used to host the website. The API that executed dot commands on SQLite3 and supplied data to webpage was implemented on Python Flask. HTML, JQuery and Bootstrap CSS were used to build the front-end of the website. DC.js and Crossfilter were used to make the dynamic chart features on the website.

The entity relationship diagram (ERD) was designed to store 3 tables representing different kind of information obtained from the analysis: GenomeDetail, QualityScore, Taxonomy. As shown in Figure 2, each entry in each of the tables demonstrates a field of information contained in the tables. The GenomeDetail table contained name and identifier of all genomes along with basic details such as genome size, number of contigs, GC content. A QualityScore table stored the genome quality scores and other 4 quality scores. A well curated taxonomy data related to bacterial genomes in GenBank were downloaded from a Namesforlife website. This taxonomy data was reduced to a non-redundant set and then assigned to each genome to make a Taxonomy table.

**Figure 2.**
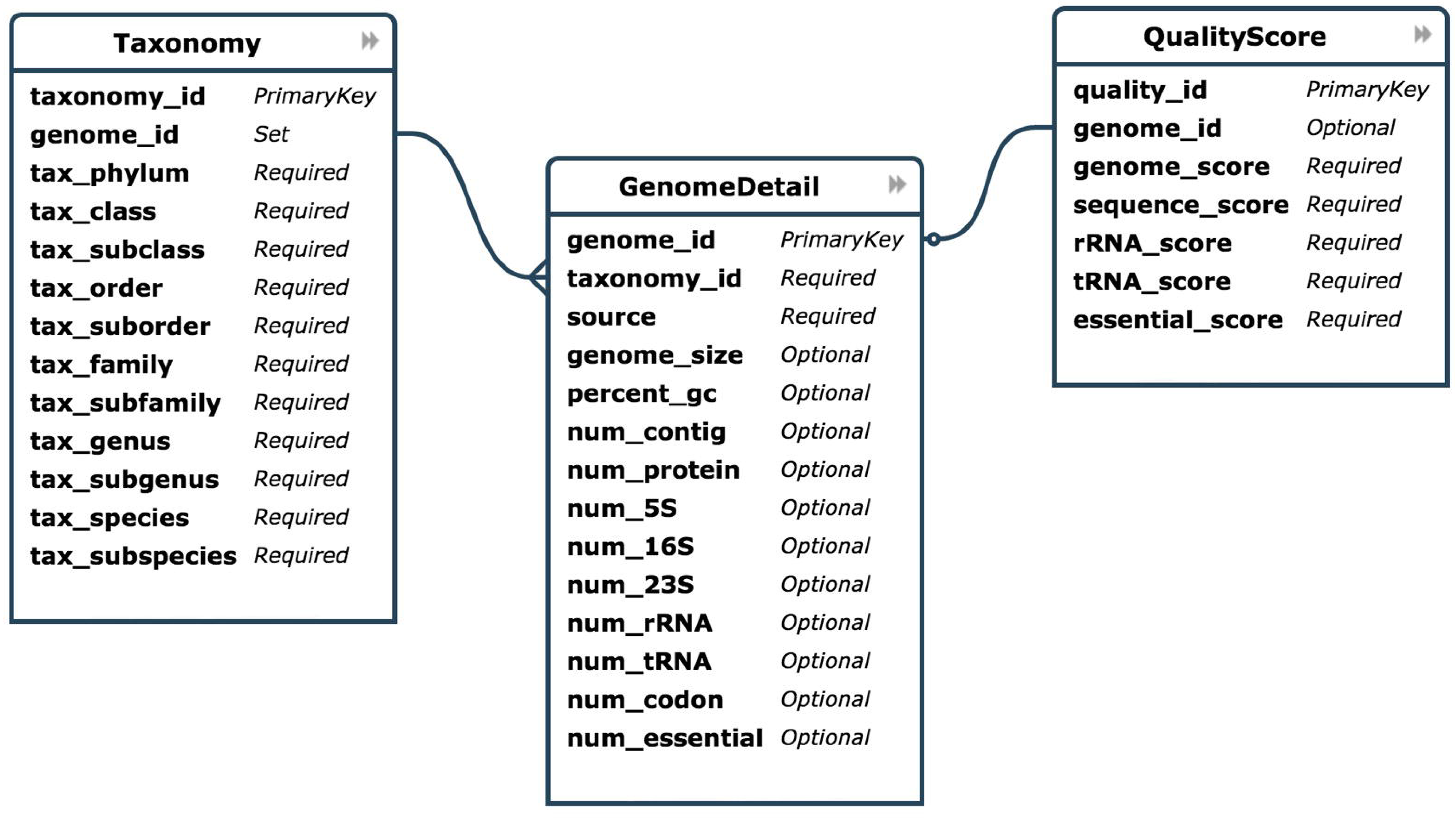
The entity relationship diagram of tables stored in the database of dBBQs.

## Utility and discussion

The total number of bacterial genomes stored in the database of dBBQs is 96,167 genomes. These genomes were collected from 4 different genome repositories: 67,980 genomes from GenBank; 11,768 genomes from GenBank - SRA; 2,477 genomes from Broad; 11,944 genomes from Kbase; 1,998 genomes from PATRIC. According to the “safe-to-use” genome quality score at 0.8 or better, we found that 65,689 out of 96,167 (∼68%) genomes passed this criterion. Table 1 shows the summary of number of bacterial genomes, genome quality scores, and 4 scores for different sources. As expected, the average of genome quality scores of 4 different sources met the safety criterion except genomes from GenBank - SRA that have the average score at 0.69. The low average genome quality score was usually because there were too many contiguous pieces for each genome which significantly brought the average sequence quality score down too low and affected the genome quality score.

**Table 1.** Number of genomes and average scores for each data source used in dBBQs.

| Data source | Number of genomes | Average of Genome Quality Score | Average of Raw Quality Scores |  |  |  |
| --- | --- | --- | --- | --- | --- | --- |
|  |  |  | Sequence Quality Score | rRNA Score | tRNA Score | Essential Gene Score |
| Genbank | 67980 | 0.85 | 0.79 | 0.69 | 0.91 | 0.99 |
| Genbank - SRA | 11768 | 0.69 | 0.38 | 0.72 | 0.74 | 0.92 |
| Broad | 2477 | 0.94 | 0.9 | 0.93 | 0.95 | 0.97 |
| Kbase | 11944 | 0.9 | 0.82 | 0.86 | 0.93 | 0.99 |
| Patric | 1998 | 0.9 | 0.81 | 0.84 | 0.96 | 0.99 |
| **Total** | 96167 | 0.856 | 0.74 | 0.808 | 0.898 | 0.972 |

Comparing the annotations between dBBQs and the original source databases remains a difficult task due to the lack of provided complete annotations for all genomes. However, we still can compare the number of predicted proteins as it is the most complete annotation in the database of bacterial genomes. For purposes of assessing quality of protein prediction, we downloaded the metadata which contains numbers of predicted proteins of all genomes from NCBI [https://www.ncbi.nlm.nih.gov/genome/browse]. As can be seen in Figure 3, we compared the distribution of predicted proteins between dBBQs and GenBank in 4 different levels of genome status (Complete Genome, Chromosome, Contig, Scaffold). dBBQs showed very similar at locating proteins in most of genomes in GenBank with a few exceptions even in scaffolds which contain lots of contigs and gaps.

**Figure 3.**
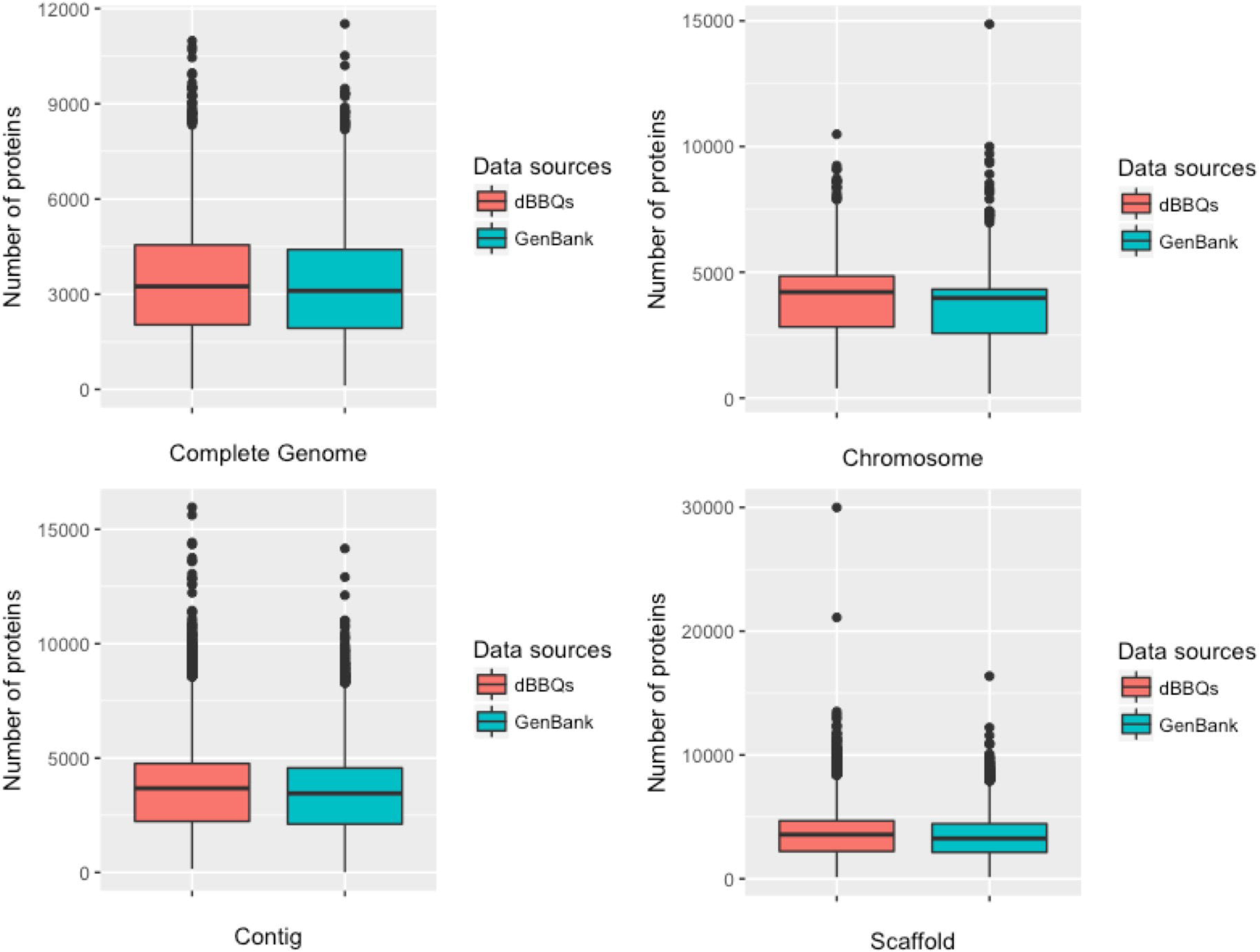
Comparison of protein prediction between dBBQs and GenBank.

## User interface

### Interactive Chart Section

Figure4 shows the front page of dBBQs which composes of 2 types of chart (6 bar charts of ‘Genome Quality score’, ‘Sequence Quality Score’, ‘rRNA Score’, ‘tRNA Score’, ‘Essential Gene Score’, and ‘Taxonomy: Phylum’; 1 donut chart of ‘Genome Repositories’) and 1 table of genome information. User can select the data category or range of scores from all charts as filters to display on the website. Once any of charts is selected, the other charts will therefore be updated instantly. For example, when GenBank is selected from the donut chart of Genome Repositories, all bar charts and table will update their information dynamically. To be easy for user to focus at the main score first, we differentiated by picking different colors. The color of bar chart of genome quality score is in red color while other scores is in blue color.

**Figure 4.**
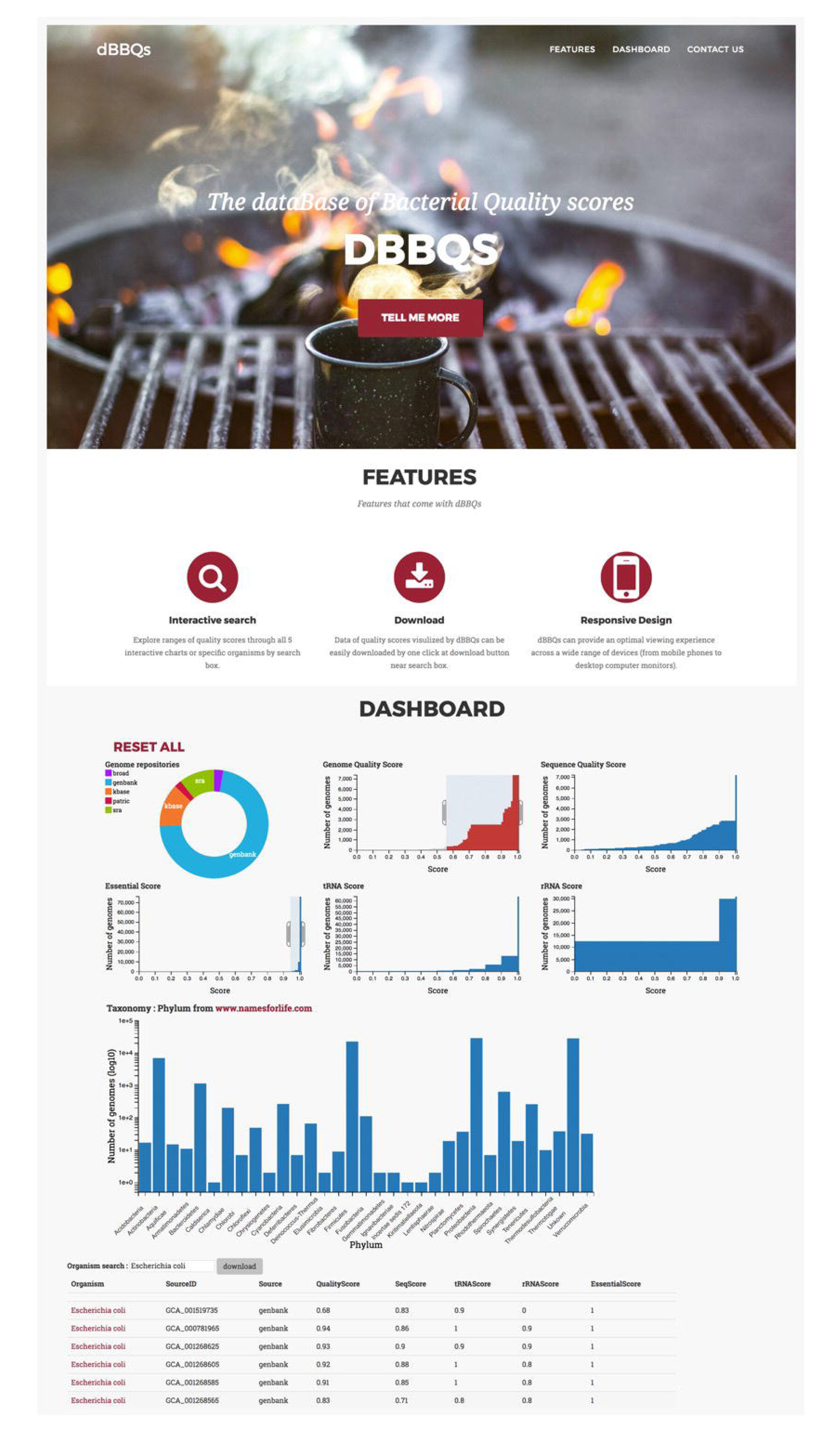
The front page of dBBQs database and interactive charts of all quality scores.

### Search Section

Through the dBBQs front page, it is integrated with the search function. It can be found at a search box below the Taxonomy bar chart. This search box allows users to scope the group of bacterial genomes by searching for the name. For example, when the ‘Escherichia coli’ search term was supplied to the search box all charts and table were filtered and displayed for only genomes containing a word ‘Escherichia coli’ in their name. Furthermore, on the right of search box, users can download the search results in the table for the further analysis. A result file will be generated in CSV format, which can be open on many spreadsheet programs such as Microsoft Excel and Number.

### Genome quality and statistics section

Any information in detail can be retrieved by clinking at the name of genome on the table. The genome quality and statistics page will start at the new tab on the browser (Figure5). This page comprises of 5 sections represented by 5 different frames: details of ‘all scores and taxonomy’ in white frame, details of ‘sequence quality score’ in green frame, details of ‘rRNA score’ in blue frame, details of ‘tRNA score’ in yellow frame, and details of ‘essential gene score’ in red frame. In addition, users can download data in 5 sections above in CSV format by clicking at the button next to the Genome Quality Score. These attributes will give users details behind a calculation of all quality scores and leverages complex analytics when genomes have very similar scores but different in some details.

**Figure 5.**
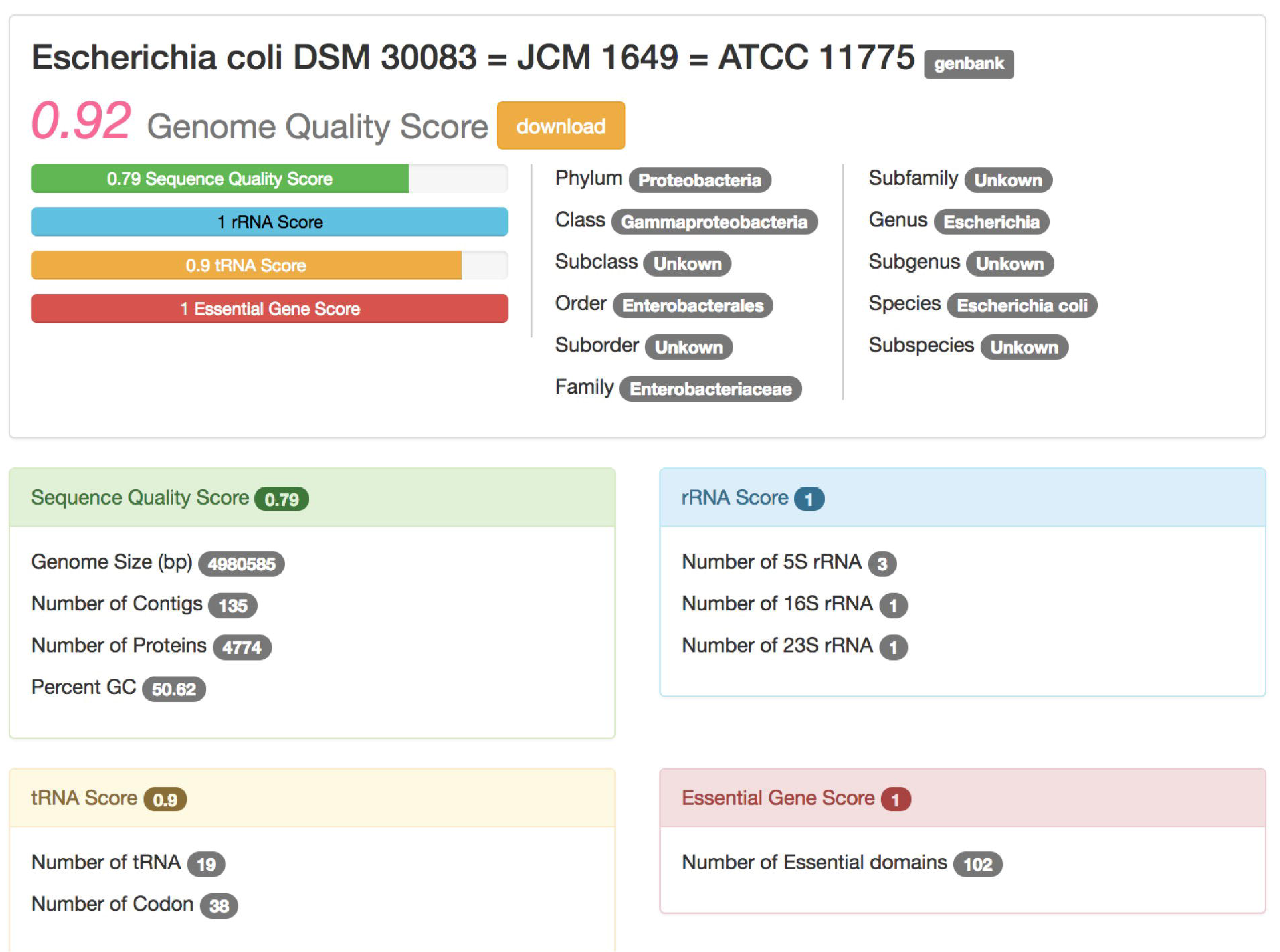
Genome quality score and statistics card for Acaricomes phytoseiuli DSM 14247.

## Conclusion

dBBQs provides quality scores for all available genome sequences with a user-friendly Web-based user interface. These scores can be used as one of cut-offs to get a high-quality set of genomes for testing bioinformatics tools or improving the analysis. Additionally, all data of four measurements that were combined to make the quality score for each genome, can be download in CSV format. The data table can be imported to a network and molecular profiling tool like CytoScape. By using CytoScape, the data table can potentially be used as node attributes for further analysis on pathway comparison using KEGG or BioCyc plugins.

Moreover, we plan to release our API to support the connection between other bioinformatics websites and our database. Also, a Web tool for calculating of quality score will be added to the website to allow users to upload genome sequences and get the genome quality scores. The database of dBBQs will be update regularly as number of genomes in public databases growing rapidly and is freely use for non-commercial purpose. These extensions of functionality and long term intention will help contribute largely to the analysis of quality of genomic data in bacterial research community.

## Availability and requirements

Project name: dBBQs: dataBase of Bacterial Quality scores

Project home page: http://arc-gem.uams.edu/dbbqs

Operation system(s): Web based, Platform independent

Programming language: HTML, CSS, JavaScript, Python

## List of abbreviations

dBBQs: dataBase of Bacterial Quality scores
NCBI: National Center for Biotechnology information
Kbase: The U.S. Department of Energy Systems Biology Knowledgebase
PATRIC: Pathosystems Resource Integration Center
SRA: Sequence Read Archive
API: Application program interface
CSV: Comma-separated Value.

## Declarations

### Authors’ contributions

VW designed the project, collected data, performed the analysis, wrote the website, and drafted the manuscript. PP contributed back-end coding and constructed the database. IN helped design the website, discussed the results and interpretation of final data. DU conceived and directed the project. All participated in finalizing and approved the manuscript.

## Acknowledgements

The authors acknowledge the Texas Advanced Computing Center (TACC) at The University of Texas at Austin, the Arkansas High Performance Computing Center (AHPCC), multiple National Science Foundation grants, and the Arkansas Economic Development Commission for providing HPC resources that have contributed to the research results reported within this paper. URL: http://www.tacc.utexas.edu and http://hpc.uark.edu.

## Ethics approval and consent to participate

Not applicable.

## Consent for publication

Not applicable.

## Availability of data and materials

The datasets generated and/or analysed during the current study are available in the dBBQs database, http://arc-gem.uams.edu/dbbqs

